# Impoverished auditory cues limit engagement of brain networks controlling spatial selective attention

**DOI:** 10.1101/533117

**Authors:** Yuqi Deng, Inyong Choi, Barbara Shinn-Cunningham, Robert Baumgartner

## Abstract

Spatial selective attention enables listeners to process a signal of interest in natural settings. However, most past studies on auditory spatial attention used impoverished spatial cues: presenting competing sounds to different ears, using only interaural differences in time (ITDs) and/or intensity (IIDs), or using non-individualized head-related transfer functions (HRTFs). Here we tested the hypothesis that impoverished spatial cues impair spatial auditory attention by only weakly engaging relevant cortical networks. Eighteen normal-hearing listeners reported the content of one of two competing syllable streams simulated at roughly +30 ° and −30° azimuth. The competing streams consisted of syllables from two different-sex talkers. Spatialization was based on natural spatial cues (individualized HRTFs), individualized IIDs, or generic ITDs. We measured behavioral performance as well as electroencephalographic markers of selective attention. Behaviorally, subjects recalled target streams most accurately with natural cues. Neurally, spatial attention significantly modulated early evoked sensory response magnitudes only for natural cues, not in conditions using only ITDs or IIDs. Consistent with this, parietal oscillatory power in the alpha band (8-14 Hz; associated with filtering out distracting events from unattended directions) showed significantly less attentional modulation with isolated spatial cues than with natural cues. Our findings support the hypothesis that spatial selective attention networks are only partially engaged by impoverished spatial auditory cues. These results not only suggest that studies using unnatural spatial cues underestimate the neural effects of spatial auditory attention, they also illustrate the importance of preserving natural spatial cues in assistive listening devices to support robust attentional control.

**Highlights:**

- Neural responses are weak or even absent with impoverished spatial auditory cues.
- Spatial cue realism affects parietal alpha activity and early evoked cortical responses.
- Differences due to cue realism disappear by the next level of neural processing.
- Robust engagement of spatial attention mechanisms requires realistic spatial cues.

## 1 Introduction

Spatial hearing is crucial to selectively attend to sounds of interest in everyday social settings. The remarkable ability of normal-hearing listeners to focus on a sound source within a complex acoustic scene is often referred to as “the cocktail party phenomenon,” and has a rich history (Cherry, 1953). Nevertheless, the mechanisms controlling spatial selective attention are still poorly understood. Acoustically, in everyday situations, the two ears provide the listener with a listener-specific combination of spatial cues that include interaural time and intensity differences (ITDs and IIDs, respectively), as well as spectral cues caused by acoustical filtering of the pinnae (Blauert, 1997a). Together, these cues, captured by individualized head-related transfer functions (HRTFs), allow the brain to create a clear, punctate internal representation of the location of sound sources in the environment (Majdak et al., 2019; Middlebrooks, 2015).

When only isolated or impoverished spatial cues are present, auditory localization performance degrades and the natural perception of external auditory objects may even collapse into the listener’s head (Baumgartner et al., 2017; Callan et al., 2013; Cubick et al., 2018; Hartmann and Wittenberg, 1996). Nevertheless, degraded or isolated ITDs and IIDs still create a strong sense of lateralization within the head; moreover, even highly impoverished spatial cues can be used to achieve spatial release from speech-on-speech masking, behaviorally (Cubick et al., 2018; Culling et al., 2004; Ellinger et al., 2017; Glyde et al., 2013; Kidd et al., 2010; Loiselle et al., 2016). The relative importance of ITDs and IIDs in spatial release from masking remains unclear, with past studies reporting conflicting results when directly comparing different binaural conditions (Ellinger et al., 2017; Glyde et al., 2013; Higgins et al., 2017; Shinn-Cunningham et al., 2005). More importantly, it is a puzzle as to why realistic and degraded spatial cues yield at best small behavioral differences in masking release even though spatial perception is clearly degraded when cues are impoverished.

Previous electroencephalography (EEG) and magnetoencephalography (MEG) studies have demonstrated that rich spatial cues in sound stimuli lead to different cortical activity compared to using isolated cues during sound localization (Callan et al., 2013; Palomäki et al., 2005) and auditory motion processing (Getzmann and Lewald, 2010). However, the apparently minor behavioral consequences of using unnatural, non-individualized spatial cues on spatial release from masking, combined with the ease of implementing studies with simple, non-individualized spatial cues, led to their wide usage in auditory neuroscience studies (Cusack et al., 2001; Dahmen et al., 2010; Dai et al., 2018; Itoh et al., 2000; Kong et al., 2014; Sach et al., 2000). Indeed, in the auditory neuroscience literature, many studies did not even present true binaural signals, but instead studied “spatial” attention by using dichotic signals, with one sound presented monaurally to one ear and a competing sound presented monaurally to the other ear (Ahveninen et al., 2011; Alho et al., 1999b; Das et al., 2016; Wöstmann et al., 2016). These studies implicitly assumed that because listeners were able to use impoverished spatial cues to listen to one sound from a particular (relative) direction, the cognitive networks responsible for controlling spatial attention must be engaged just as they are when listening to rich, natural spatial cues. Nonetheless, it is unclear whether and how engagement of higher-order cognitive processes such as deployment of selective attention is affected by the use of unnatural or impoverished spatial cues.

Modulation of neural signatures, such as event-related potentials (ERPs) and induced oscillatory activity, is often taken as evidence of effective attentional control (Herrmann and Knight, 2001; Siegel et al., 2012). In particular, auditory spatial attention is known to modulate early sensory ERPs in the N1 time range (processing latencies of 100 to 150 ms; see Choi et al., 2013; Röder et al., 1999), whereas modulation of P1 ERPs (50 to 100 ms) has only recently been demonstrated in a free field experiment (Giuliano et al., 2014). Induced alpha oscillation (8 to 14 Hz) has been hypothesized to function as an information gating mechanism (Klimesch et al., 2007). During auditory spatial attention, parietal alpha power often decreases in the contralateral hemisphere of attended stimuli and/or increases in the ipsilateral hemisphere (Banerjee et al., 2011; Lim et al., 2015; Wöstmann et al., 2016). These neural modulations constitute objective metrics of the efficacy of attentional control.

Here, we test listeners in a selective attention paradigm with simultaneous, spatially separated talkers. We use the aforementioned EEG measures to compare both perceptual ability and the neural signatures of attentional control for simulations with impoverished vs. natural spatial cues. Eighteen subjects performed an auditory spatial attention task with two competing streams located at roughly +30 ° and −30° azimuth (Figure 1). On every trial, listeners were cued by an auditory cue to attend to either the left or right stream and report the content of the cued stream. The competing streams consisted of syllables (/ba/, /da/ or /ga/) from two different-sex talkers. Sound stimuli (including the cuing sound) were spatialized using three different levels of naturalness and richness: 1) generic ITDs only, 2) individualized IIDs, or 3) individualized HRTFs containing all of the naturally occurring spatial cues a listener experiences in the everyday world. We show that behavioral performance is better when listeners hear natural, individualized spatial cues than when they hear impoverished cues. Importantly, only natural spatial cues yield significant attentional modulation of P1 amplitudes. Moreover, induced alpha activity is less robust and poorly lateralized with isolated spatial cues compared to rich, natural spatial cues.

**Fig. 1.**
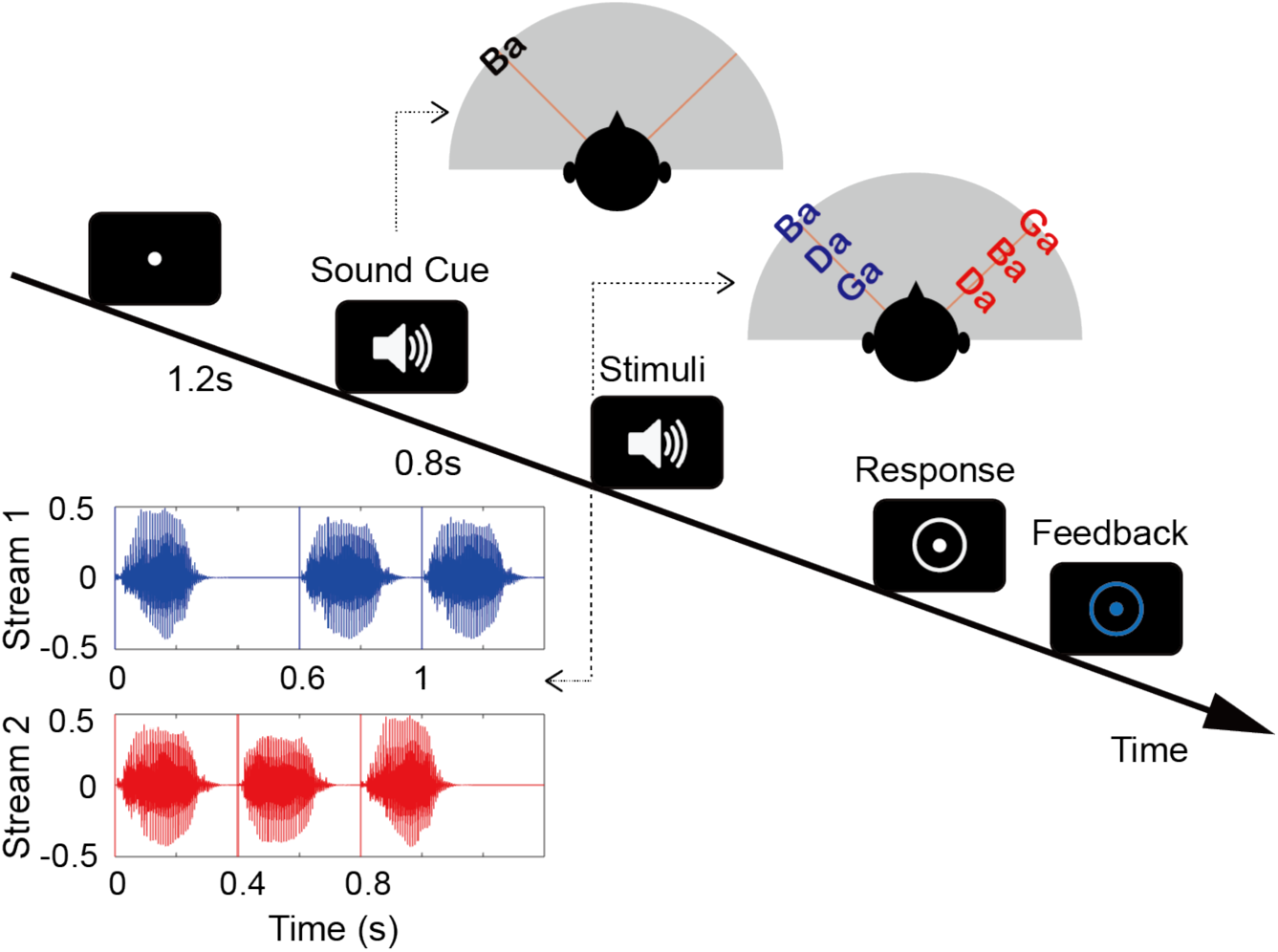
Auditory spatial attention task with two competing streams was used to assess the consequence of impoverished auditory spatial cues on neural proxies of attention control. An auditory cue was presented first from the location of the upcoming target stream, processed by the same spatialization scheme as the upcoming mixture. Following the cue, the competing streams began, one from around +30° the other from around −30° azimuth. Listeners were asked to recall the syllable sequence presented from the cued side. The first syllables of both streams were temporally aligned; however, the latter two syllables in the competing streams were staggered, enabling us to isolate neural responses to each. Feedback was provided after every trial.

## 2 Materials and Methods

### 2.1 Subjects

Twenty-one paid volunteers and one author within the age of 18-42 years (M = 22.9, SD = 5.5; 12 females, 10 males) participated in this study. None of the subjects had audiometric thresholds greater than 20 dB for frequencies from 250 Hz to 8 kHz. All participants gave informed consent as approved by the Boston University Institutional Review Board. Two subjects were withdrawn from the study due to the inability to perform the task (percentage of correct response less than 30% after training), and two subjects were removed during EEG data preprocessing due to excessive artifacts. Therefore 18 subjects remained for further analysis (N = 18).

### 2.2 Stimuli and Procedure

The sound stimuli consisted of consonant-vowel syllables (/ba/, /da/, & /ga/), each 0.4 s in duration. These syllables were recorded from three talkers that naturally differed in fundamental frequency (F0). Details on stimulus are provided in Stimulus Presentation. Cue and stimuli were presented via earphones (ER-2, Etymotic Research, Inc.) and spatialized to approximately ±30° azimuth (0° elevation). Three different spatialization conditions were used: HRTF, IID, and ITD. In the HRTF condition, individualized HRTFs, providing natural combinations of ITDs, IIDs, and spectral cues, were used.

Individualized HRTFs were measured using procedures identical to those described in a previous study (Baumgartner et al., 2017). In short, loudspeakers were positioned at the desired angles and 1.5 m distance from the subject’s head in a sound-treated chamber. A pair of miniature microphones placed at the entrances of the subject’s blocked left and right ear canals measured the pseudo noise signal emitted by each loudspeaker. These measurements were used to compute the impulse responses of the acoustic transmission paths. Room reflections were removed via temporal windowing (0.5-ms cosine ramps) limiting the impulse responses to the initial 3 ms. Finally, those listener-specific impulse responses were equalized by reference measurements obtained by placing the microphones at the radial center of the loudspeaker setup.

In the IID condition, ITDs were removed from the individualized HRTFs by computing minimum-phase representations of the filters (computed by removing the non-causal part of the cepstrum). Hence, the IID and HRTF conditions provided the same monaural magnitude spectra and thus the same energetic advantage of the ear ipsilateral to the target, although the IID condition removed the naturally occurring group delay between the signals at the two ears present in the individualized HRTFs. In the ITD condition, spatialization was based on simply delaying the signal presented to the contralateral ear by 300 µs (roughly the magnitude of the ITD present in the natural HRTFs for the sources used), thus providing no energetic advantage to the ipsilateral ear or spectral cues present in the natural HRTFs. This spatialization method was tested due to its popularity in auditory neuroscience.

The auditory cue was a single syllable /ba/ spoken by a low-pitched male voice (F0 = 91 Hz, estimated by Praat software; Boersma, 2001). The subsequent target and distractor streams each consisted of three syllables randomly chosen out of the set of three syllables (with replacement). The target stream was spoken by either a female (F0 = 189 Hz) or a high-pitched male talker (F0 = 125 Hz), and the distractor stream was spoken by the other talker. The first syllable of the target and distractor sound overlapped in time, while the latter two syllables were separated by 200 ms, onset to onset (Figure 1). To avoid engagement of temporal attention rather than spatial attention, the target stream was equally likely to be leading or lagging, randomly chosen on each trial. In the leading stream, the onsets of all three syllables were separated by 400 ms; in the lagging stream, the onsets of the first and the second syllable were separated by 600 ms, whereas those of the second and the third syllable were separated by 400 ms. All sound stimuli were presented at a sound pressure level of approximately 75 dB.

### 2.3 Task

Subjects performed a spatial attention task using a Posner paradigm (Figure 1) (Posner et al., 1980) while listening to sounds over headphones in a sound-treated booth (Eckel Industries, Inc.). Sound spatialization was realized by one of the three spatialization conditions fixed within trials but pseudo-randomized across trials. Subjects were instructed to fixate on a dot at the center of the screen at the beginning of each trial. The fixation dot lasted 1.2 s before an auditory cue was presented. The auditory cue came from either left or right, indicating the direction from which the target sound would come. A target sound started 0.8 s later from the cued location. At the same time a distractor sound started from the opposite location of the target sound. After the sounds finished, a response cue appeared on the computer screen, signaling to the subjects to report the syllable sequence of the target sound using a number keypad. The syllables /ba/, /da/ and /ga/ corresponded to number keys 1, 2, and 3, respectively. The keys were labelled with their corresponding syllables. Feedback about whether or not the subject correctly reported the syllables was given at the end of every trial.

Each subject performed 450 randomized trials of this task, divided into 9 blocks each consisting of 50 trials. In total, every subject performed 150 trials for each of the three sound spatialization conditions (75 trials attending left and 75 trials attending right; half target leading and half target lagging). Prior to the test sessions, all participants received a practice session to get familiarized with the task. Participants with a percentage of correct response below 30% after 3 blocks of training (50 trials per block) were excluded from the study.

### 2.4 EEG Acquisition and Preprocessing

32-channel scalp EEG data was recorded (Activetwo system with Activeview acquisition software, Biosemi B.V.) while subjects were performing the task. Two additional reference electrodes were placed on the earlobes. Horizontal eye movements were recorded by two electrooculography (EOG) electrodes placed on the outer canthi of each eye. Vertical eye movement was recorded by one EOG electrode placed below the right eye. The timing of stimulus was controlled by Matlab (Mathworks) with Psychtoolbox (extension 3; Brainard, 1997).

EEG preprocessing was conducted in Matlab with Eeglab toolbox (Delorme and Makeig, 2004). EEG data were corrected against the average of the two reference channels. Bad channels were marked by manual selection during recording and automatically detected based on joint probability measures of Eeglab. EEG signals were then down-sampled to 256 Hz and epochs containing responses to individual trials were extracted. Each epoch was baseline corrected against 100 ms prior to the cue onset by removing the mean of the baseline period from the whole trial. ICA artifact rejection was performed with Eeglab to remove components of eye movements, blinks, and muscle artifacts. The maximum number of independent components rejected for each subject was five. After ICA rejection, bad channels were removed and interpolated. Trials with a maximum absolute value over 80 µV were rejected (Delorme et al., 2007). Two subjects with excessive artifacts were removed from further EEG analysis because less than 50% of trials remained after thresholding. For the rest of the 18 subjects, at least about two thirds of the trials (minimum was 48 out of 75 trials) remained for each condition after artifact rejection. Trial numbers were equalized within and across subjects by randomly selecting the minimum number of available trials (N = 48) for each condition across the whole recording session.

### 2.5 Data analysis

Behavioral performance was quantified by the percentage of correct responses for each one of the three syllables in the target stream and each spatialization condition. Behavioral results were collapsed across the attend-left and attend-right trials. The percentages of correct response were then normalized by logit transformation before parametric statistical testing was performed on the resulting data.

ERP responses were evaluated for the second syllable of the target sound and distractor sound, respectively. The reason we looked at the second syllable only is that 1) the first syllable of the target and distractor aligned in time and therefore the ERPs were not separable, and 2) the ERP amplitude in response to the third syllable was small, and therefore more contaminated by noise. ERP components were then extracted from the time series data. The preprocessed data (details see EEG Preprocessing Procedures) was bandpass filtered from 0.5 to 20 Hz by a finite impulse response filter with Kaiser window design (β = 7.2, n = 1178). Data from four fronto-central channels (Cz, Fz, FC1, and FC2) were averaged to get the auditory ERP response. We picked these four channels a priori because auditory ERP responses in sensor space are largest in the fronto-central area of the scalp. To quantify the amplitudes of ERP components, the maximum value within the window of 50 to 100 ms after the second syllable onset was taken to be the P1 amplitude; the minimum value within the window of 100 to 180 ms after the second syllable onset was calculated to be the N1 amplitude. The values extracted from the selected windows were calculated for each channel and plotted onto a 2D scalp map to generate topography plots. The values of the ERP components from the four selected channels were then averaged and compared across different spatialization conditions.

To get the amplitude of alpha oscillations, the preprocessed EEG data was bandpass filtered to the alpha range (8 to 14 Hz) before a Hilbert transform was applied. The magnitude of the resulting data was taken as the extracted alpha power envelope. To get induced alpha power, the alpha power was calculated for single trials first and then averaged across trials (Snyder and Large, 2005). The time course of alpha power was baseline corrected against 700 ms before the auditory cue onset. GFP (Murray et al., 2008; Skrandies, 1990) constitutes the spatial standard deviation across all scalp electrodes; it has been used as a measurement to quantify the amount of alpha variation across the scalp (Lim et al., 2015). We calculated the time courses of alpha GFP by taking the standard deviation of alpha power over all electrodes. To quantify the degree of alpha modulation based on direction of attention, we calculated the Attentional Modulation Index (AMI) of alpha power, defined as the alpha power difference between attended left and attended right trials divided by the overall alpha power (Wöstmann et al., 2016). The AMI of alpha was calculated for each time point, yielding the time course of AMI for each spatialization condition. We then averaged the alpha AMI of each spatialization condition over the 800 ms immediately before stimulus onset (−800 ms to 0 ms, re: onset). This is the period in which the cue has already signaled to the subjects where to orient their spatial attention in preparation for the target sound, but before the speech streams begin. Scalp topographies of the preparatory alpha AMI were plotted for each condition. Hemispheric lateralization of alpha AMI was further compared across spatialization conditions and evaluated as the difference between the left hemisphere and the right hemisphere. Calculated in this way, the AMI is expected to be positive in left and negative in right parietal channels.

For testing the significance of different means across conditions, we conducted repeated measures ANOVAs followed by post-hoc analyses for all significant main effects and interactions using Fisher’s least significant difference procedure. We separately tested whether condition means differed significantly from zero using Bonferroni-corrected t-tests (*P*_*adj*_). The Lilliefors test was performed prior to statistical testing to check normality of the data. Data was considered normally distributed at *P* > 0.05. Prior to statistical analysis of behavioral performance, the percentages of correctly reported syllable were logit transformed in order to obtain normally distributed data.

Raw data and analysis scripts are publicly available (Deng et al., 2019).

## 3 Results

### 3.1 Natural spatial cues facilitate behavioral performance

Percentages of correctly recalling each syllable of the target stream differed across the three spatialization conditions (Figure 2; 1^st^ syllable: *F*_(2,34)_ = 25.25, *P* < 0.001; 2^nd^ syllable: *F*_(2,34)_ = 6.27, *P* = 0.005; 3^rd^ syllable: *F*_(2,34)_ = 5.60, *P* = 0.008). For the first syllable, where the target and distractor sounds overlapped in time, subjects were least accurate in the ITD condition; performance in the ITD condition differed significantly from both the IID (t_(34)_ = 5.31, *P* < 0.001) and HRTF conditions (t_(34)_ = 6.74, *P* < 0.001). However, no statistically significant difference was observed between IID and HRTF conditions for that syllable (t_(34)_ = 1.43, *P* = 0.16). For the second and the third syllable, where target and distractor streams occurred staggered in time, subjects performed significantly better in the HRTF condition than in both the ITD condition (2^nd^ syllable: t_(34)_ = 3.27, *P* = 0.002; 3^rd^ syllable: t_(34)_ = 3.33, *P* = 0.002) and the IID condition (2^nd^ syllable: t_(34)_ = 2.81, *P* = 0.008; 3^rd^ syllable: t_(34)_ = 1.94, *P* = 0.06). There was no significant difference in performance between the ITD and IID conditions for the two staggered syllables (2^nd^ syllable: t_(34)_ = 1.41, *P* = 0.17; 3^rd^ syllable: t_(34)_ = 1.39, *P* = 0.17).

**Fig. 2.**
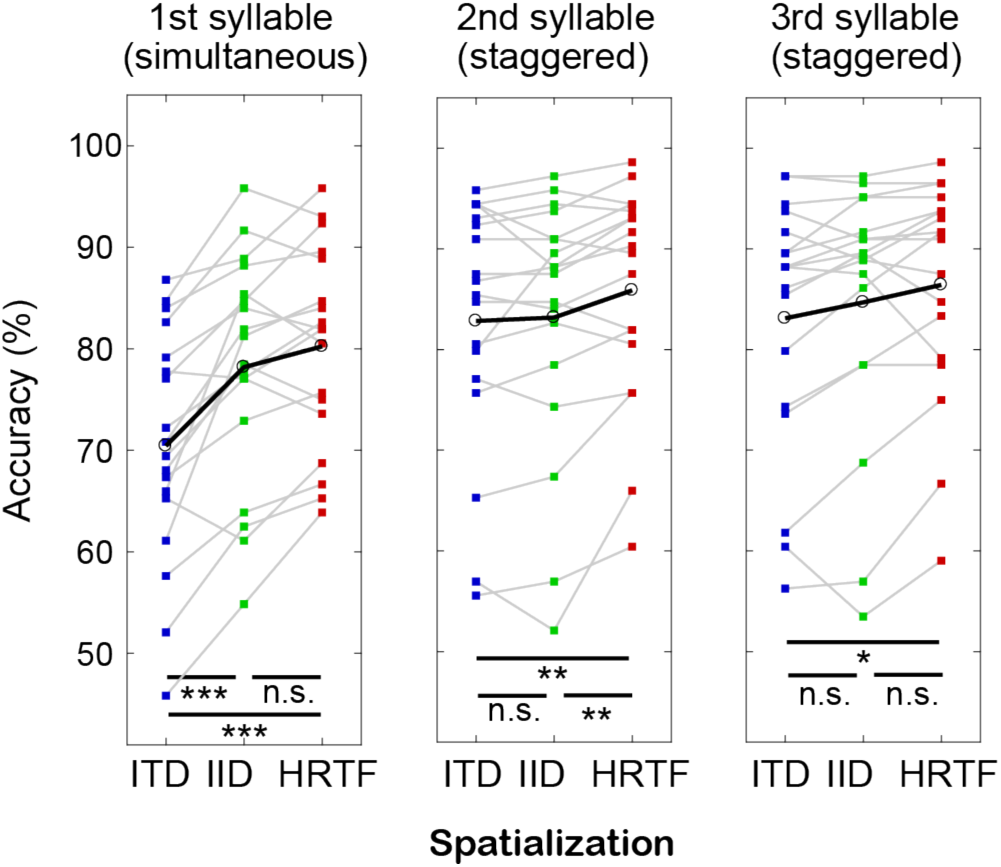
Listeners’ (N = 18) recall performance was evaluated for every syllable and every spatialization condition. Sounds were spatialized either based on generic ITDs, individualized IIDs, or the natural combination of ITDs, IIDs, and spectral cues in individualized HRTFs. Behavioral advantages of having more consistent spatial information were statistically significant but small in absolute terms. * P < .05; ** P < .001; *** P < .0001

### 3.2 Impoverished spatial cues affect attentional modulation of ERPs

Figure 3A shows the ERPs evoked by the onset of the second syllable of the attended target sound and the unattended distractor sound, aligning the onsets of the target and distractor syllables to 0 s to allow direct comparison. Stimulus onsets elicited a fronto-central positivity (P1) between 50 to 100 ms followed by a negativity (N1) between 100 to 180 ms (Figure 3A-B). The amplitudes of these two components were extracted and the difference between attended stimuli (target sound) and unattended stimuli (distractor sound) was calculated in order to quantify attentional modulation for both the P1 and N1 components (Figure 3C).

**Fig. 3.**
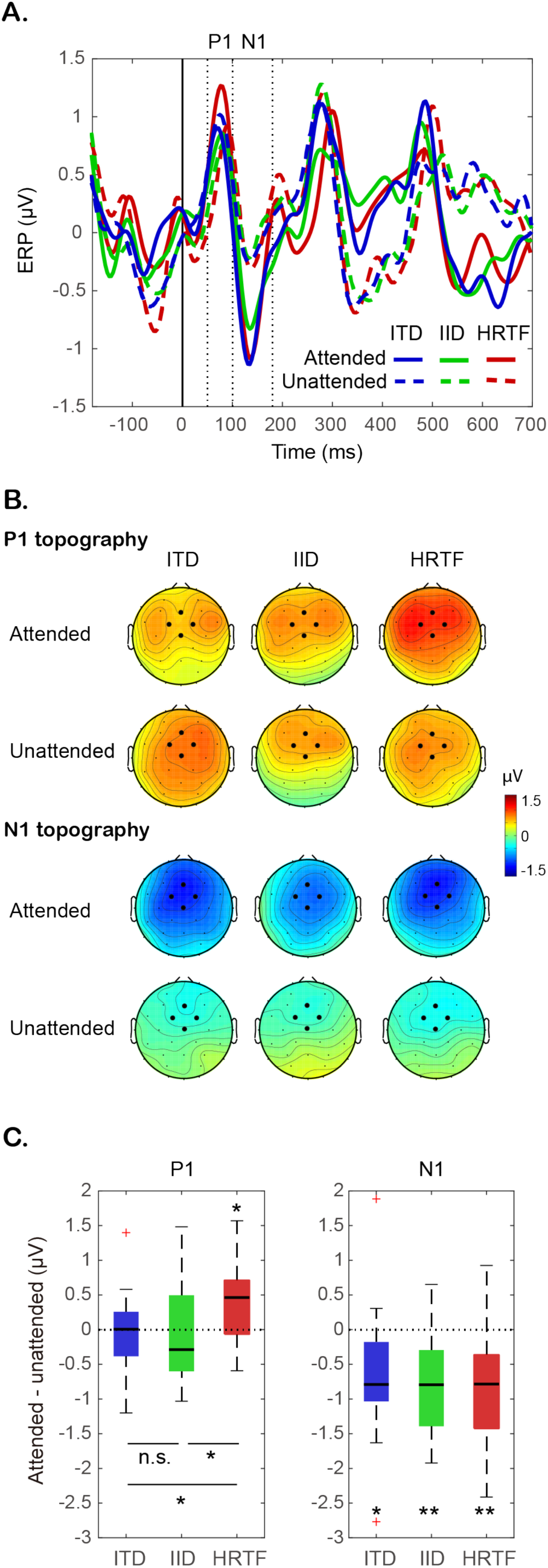
P1 amplitudes were only modulated by the attended direction in the HRTF condition, whereas N1 amplitudes were modulated equally strongly across spatialization conditions (N = 18). **A.** ERP waveforms at fronto-central electrodes were compared between the attended target stream and the unattended distractor stream for every spatialization condition. The P1 time range was defined as 50 ms to 100 ms, and the N1 time range as 100 ms to 180 ms. **B.** Most topographies of both ERP components show maxima at the fronto-central sites (black dots) used for evaluation. C. The modulation strength of ERP components was assessed by the amplitude differences between attended and unattended streams. * P < .05; ** P < .01

We tested whether P1 responses were significantly larger to attended stimuli than to unattended stimuli separately for each of the three spatialization conditions. Only the HRTF condition showed a significant P1 modulation (t_(17)_ = 3.12, *P*_*adj*_ = 0.017); no significant attentional modulation was found in either the ITD (t_(17)_ = 0.50, *P*_*adj*_ = 1) or IID conditions (t_(17)_ = 0.06, *P*_*adj*_ = 1). Across conditions we found a statistically significant main effect of spatial cue on P1 amplitude modulation (*F*_(2,34)_ = 3.34, *P* = 0.047). Post hoc tests showed that attentional modulation was significantly larger in the HRTF condition than in the ITD (t_(34)_ = 2.38, *P* = 0.023) and IID conditions (t_(34)_ = 2.07, *P* = 0.046); however, modulation did not differ significantly between the ITD and IID conditions (t_(34)_ = 0.31, *P* = 0.76) (Figure 3C).

In all three spatialization conditions, the N1 amplitude was modulated significantly by spatial attention, that is, attended sounds evoked larger N1 amplitudes than unattended sounds (ITD: t_(17)_ = 3.01, *P*_*adj*_ = 0.024; IID: t_(17)_ = 4.12, *P*_*adj*_ = 0.002; HRTF: t_(17)_ = 3.56, *P*_*adj*_ = 0.007). Across the three spatialization conditions the magnitude of N1 modulation did not differ significantly (*F*_(2,34)_ = 0.060, *P* = 0.94; Figure 3C).

### 3.3 Alpha oscillation power shows less attentional modulation with impoverished spatial cues

To investigate the effect of spatialization on attentional control, we analyzed the power in alpha oscillations during the attentional preparation period (−800 ms to 0 ms), a time period in which listeners knew where to orient spatial attention based on the preceding acoustic cue, but before the sound mixture of competing streams began. We averaged the power in alpha across all trials for each spatialization condition, regardless of where spatial attention was focused, to get a measure of the total engagement of alpha activity. We then compared relative power for different attentional directions. On average across directions of attentional focus, we calculated the time courses of alpha global field power (GFP, Figure 4A) and compared within-subject differences of the temporal average within the preparatory time period across spatialization conditions (Figure 4B).

**Fig. 4.**
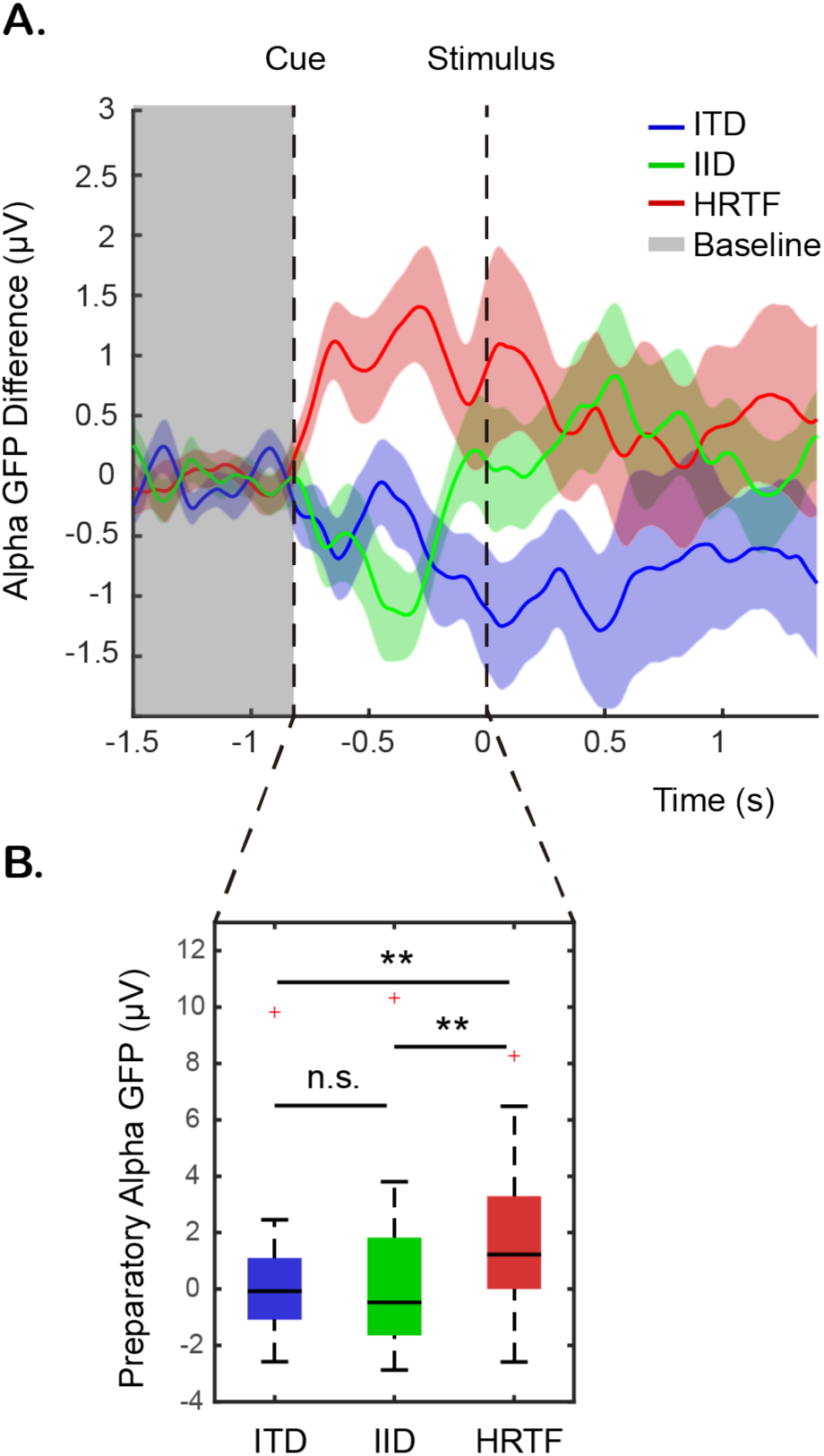
Within-subject differences in alpha-band GFP are larger in the HRTF condition, especially during the preparatory time window (after the sound cue but before the first syllables of the competing streams). **A.** Waveforms of the average (± SEM) GFP differences are shown during the baseline period, preparatory phase, and stimulus phase with stream competition. **B.** The temporal average of the preparatory alpha GFP difference is larger for the HRTF condition. ** P < .01

Alpha GFP was not significantly modulated in either the ITD or IID conditions (ITD: t_(17)_ = 0.44, *P*_*adj*_ = 1; IID: t_(17)_ = 0.43, *P*_*adj*_ = 1), while in the HRTF condition, the GFP tended to be greater than zero (HRTF: t_(17)_ = 2.56, *P*_*adj*_ = 0.061). In a direct comparison, spatialization conditions differed significantly in alpha GFP (*F*_(2,34)_ = 5.26, *P* = 0.010). In particular, alpha GFP in the HRTF condition was significantly larger than in each of the other two conditions (HRTF vs ITD: t_(34)_ = 2.80, *P* = 0.008; HRTF vs IID: t_(34)_ = 2.82, *P* = 0.008). No significant difference was found between the ITD and IID conditions (t_(34)_ = 0.019, *P* = 0.99).

We next assessed the lateralization of alpha power with the spatial focus of attention by comparing AMI differences across hemispheres (Figure 5). In general, the scalp topographies of AMIs show the expected hemispheric differences. However, statistically significant hemispheric differences were found only in the HRTF condition (t_(17)_ = 3.09, *P*_*adj*_ = 0.020), not in either the ITD (t_(17)_ = 1.29, *P*_*adj*_ = 0.64) or the IID condition (t_(17)_ = 0.15, *P*_*adj*_ = 1). A direct comparison of these hemisphere differences across conditions revealed a trend in which the HRTF condition had larger differences in AMI across hemispheres (*F*_(2,34)_ = 2.98, *P* = 0.064).

**Fig. 5.**
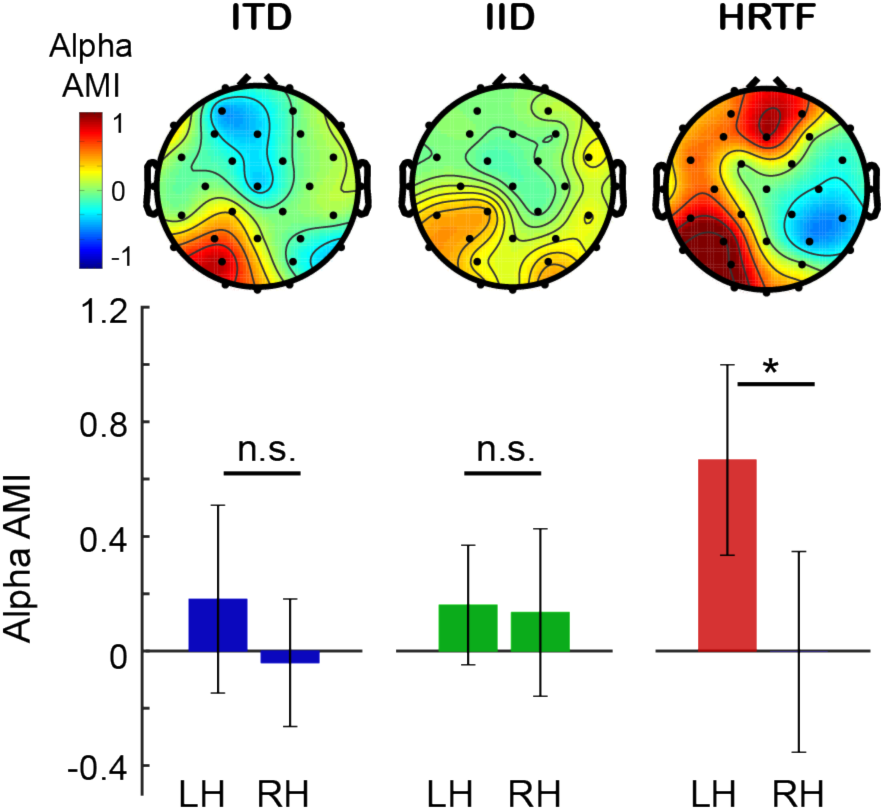
Attentional modulation of alpha activity was lateralized to the hemisphere ipsilateral to the target stream only in the HRTF condition. AMI topographies and hemispheric averages are shown for every spatialization condition (N = 18). * P < .05

In summary, impoverished spatial cues lead to worse behavioral performance, smaller P1 modulation, reduced modulation of preparatory alpha power GFP, and reduced lateralization of alpha power with attentional focus, confirming our hypothesis that impoverished spatial cues impaired engagement of spatial attention.

### 3.4 Relationships between Attentional Modulation Metrics

Given these consistent effects of spatialization on performance and neural metrics, we explored, post hoc, whether there were ordered relationships in the individual measures of attentional control, including P1 modulation, preparatory alpha GFP, and alpha power lateralization. To investigate the relationship between evoked response modulation and alpha oscillatory activity, we first calculated the regression slope relating P1 amplitude to preparatory alpha GFP for each subject, and then performed a paired t-test on the coefficients obtained. No consistent relationship between alpha GFP and P1 amplitudes was observed (t_(17)_ = 0.90, *P* = 0.38). Correlation analysis was also conducted comparing behavioral accuracy to P1 modulation, defined as the attended P1 amplitude minus unattended P1 amplitude. No consistent relationships between P1 modulation and behavioral performance were observed for any syllable (1st syllable: t_(17)_ = 0.54, *P* = 0.59; 2nd syllable: t_(17)_ = 0.31, *P* = 0.76; 3rd syllable: t_(17)_ = 0.69, *P* = 0.50). Similarly, we did not observe consistent relationships between alpha AMI lateralization and response accuracy for any syllable (1st syllable: t_(17)_ = 0.19, *P* = 0.85; 2nd syllable: t_(17)_ = 1.39, *P* = 0.18; 3rd syllable: t_(17)_ = 0.11, *P* = 0.91). In addition, no consistent relationship was found between alpha GFP and response accuracy for any syllable (1st syllable: t_(17)_ = 0.65, *P* = 0.52; 2nd syllable: t_(17)_ = 1.27, *P* = 0.22; 3rd syllable: t_(17)_ = 1.16, *P* = 0.26). Thus, although there were significant differences in engagement of attention across spatial conditions as measured both behaviorally and neurally, the individual subject differences in these metrics were not closely related.

## 4 Discussion

Behaviorally, we found that impoverished spatial cues impair performance on an auditory spatial attention task in a multi-talker scene. We used objective electrophysiological measures to assess whether the naturalness and richness of spatial cues also impacts how strongly auditory spatial attention modulates brain responses. We found that impoverished spatial cues reduce the strength of the evoked and induced neural signatures of attentional control. Specifically, evoked P1 amplitudes and induced alpha oscillatory power showed less attentional modulation for sound stimuli with impoverished spatial cues compared to when spatial cues were tailored to recreate the natural, rich experience of individual listeners.

### 4.1 Impoverished spatial cues result in less neural modulation during selective attention

We investigated attentional modulation of four established neural signatures of selective attention: evoked P1 and N1 amplitudes and induced power and lateralization of alpha oscillation. While attentional modulation of N1 amplitude was observed in all conditions, attentional modulation of the earlier P1 amplitude was not observed or was significantly weaker in the impoverished cue conditions compared to the natural cue condition. Similarly, we found less preparatory alpha power activity in the impoverished spatial cue conditions than in the natural cue condition, reflected by two indexes quantifying the amount of spatial variability of alpha power: alpha GFP (Figure 4) and AMI (Figure 5). In the ITD and IID conditions, although there was a hint of preparatory alpha lateralization over parietal sensors, the amount of lateralization was significantly smaller than in the HRTF condition and did not reach statistical significance.

Preparatory alpha activity during spatial attention tasks has been well documented to form a specific lateralization pattern in both vision and audition (Banerjee et al., 2011; Kelly, 2006; Sauseng et al., 2005; Worden et al., 2018), which is thought to be evidence of a preparatory information-gating mechanism (Foxe and Snyder, 2011; Jensen and Mazaheri, 2010; Klimesch, 2012; Klimesch et al., 2007). In vision, alpha lateralization has been observed to increase with the laterality of attention focus (Rihs et al., 2007; Samaha et al., 2015), reflecting an inhibition pattern topographically specific to attention focus.

Moreover, evidence for active top-down control of the phase of alpha oscillation during visual spatial attention suggests that alpha oscillatory activity represents active engagement and disengagement of the attentional network (Samaha et al., 2016). In addition, a previous somatosensory study revealed that the alpha lateralization is positively correlated to pre-stimulus cue reliability, further suggesting that alpha lateralization reflects top-down control that optimizes the processing of upcoming stimuli (Haegens et al., 2011). Although relatively few studies have investigated alpha activity in audition, studies suggest that alpha control mechanisms are supra-modal rather than sensory specific (Banerjee et al., 2011).

In the current experiment, a pre-stimulus auditory cue directed listeners where to focus attention in an upcoming sound mixture. The cue was spatialized using the same auditory features used to spatialize the stream mixture. Our results thus suggest that compared to stimuli with natural spatial cues, stimuli featuring only ITDs or only IIDs are less reliable in directing attentional focus, producing weaker engagement of spatial attention and reduced attentional modulation of neural responses.

Consistent with the idea that impoverished spatial cues lead to weaker engagement of spatial attention, we found that the P1 ERP component was modulated by attention only with natural spatial cues, not with impoverished cues; this result is consistent with a weak spatial representation failing to engage attentional modulation of early sensory responses (Figure 3). Our finding that attentional focus leads to a modulation of P1 amplitude for natural spatial cues is consistent with reported effects of attention on the P1 amplitude observed in previous spatial attention studies across sensory modalities [auditory: (Giuliano et al., 2014); visual: (Hillyard and Anllo-Vento, 1998; Hopfinger et al., 2004)]. Past studies agree that P1 modulation reflects an early sensory inhibition mechanism related to suppression of task-irrelevant stimuli. Although debates remain as to whether P1 modulation results from bottom-up sensory gain control (Hillyard and Anllo-Vento, 1998; Luck, 1995; Slagter et al., 2016) or some top-down inhibitory process (Freunberger et al., 2008; Klimesch, 2011), it is generally accepted in visual spatial studies that greater P1 amplitude modulation is associated with greater inhibition of to-be-ignored stimuli (Couperus and Mangun, 2010; Hillyard and Anllo-Vento, 1998; Klimesch, 2012).

Interestingly, attentional modulation of auditory P1 has been found to be positively correlated with visual working memory capacity, a result that was used to argue that stronger P1 modulation reflects better attentional control of the flow of sensory information into working memory (Fukuda and Vogel, 2009; Giuliano et al., 2014). Our result is consistent with the hypothesis that P1 modulation directly reflects attentional control. Specifically, impoverished spatial cues likely produce a “muddy” representation of auditory space that supports only imprecise, poorly focused top-down spatial attention. The resulting lack of control and specificity of spatial auditory attention results in early P1 responses that are unmodulated by attentional focus.

N1 modulation is well documented as a neural index of attentional control (Choi et al., 2013; Hillyard et al., 1998; Stevens et al., 2008; Wyart et al., 2012). The attentional modulation of N1 is thought to reflect attentional facilitation rather than inhibition (Couperus and Mangun, 2010; Marzecová et al., 2018; Slagter et al., 2016). In contrast to preparatory alpha and P1, we found that the later N1 evoked response was modulated similarly, regardless of the richness and naturalness of spatial cues.

Due to the robustness and relatively large amount of modulation, changes in auditory N1 amplitude have been used as a biomarker and a primary feature for classification of attentional focus (Blankertz et al., 2011; Schreuder et al., 2011); see also recent work on decoding attentional focus for running speech using the correlation between neural responses and the power envelope of the speech streams: (Chait et al., 2010; Mesgarani and Chang, 2012; Rimmele et al., 2015). However, there is little known about how N1 amplitudes reflect the processing of different spatial cues during auditory spatial attention. Previous studies have revealed different N1 topographies during ITD and IID processing, leading to the conclusion that ITD and IID are processed by different neural populations in the auditory cortex (Johnson and Hautus, 2010; Tardif et al., 2006; Ungan et al., 2001). However, debates remain about whether this difference in topography depends on perceived laterality, instead of different neural populations specialized for processing different spatial cues. Results from a more recent study show that auditory N1 modulation does not differ across spatial cue conditions, indicating integrated processing of sound locations in auditory cortex regardless of cues (Salminen et al., 2015). In the current study, N1 modulation did not differ across the three spatialization conditions. Thus, our results support the idea that the same cortical neural population is responsible for processing different binaural spatial cues.

### 4.2 Behavioral disadvantages associated with impoverished spatial cues are modest and depend on sound stimulus characteristics

Despite the influence of spatial cue richness on neural metrics, our behavioral results showed only small (albeit significant) behavioral differences between impoverished spatial cues and natural, individualized spatial cues (Figure 2). In line with previous studies that observed greater spatial release from masking with combined spatial cues compared to with isolated cues (Culling et al., 2004; Ellinger et al., 2017), accuracy was best in the HRTF condition. The small accuracy improvement over using impoverished cues is seen consistently across subjects. In the first syllable where the target and distractor streams overlap in time, the HRTF condition yielded a 13% increase in accuracy over the ITD condition, but is comparable to performance in the IID condition. In the two staggered syllables, accuracy in the HRTF condition is greater than in the ITD and IID conditions by only about 6% and 1%, respectively. These differences in behavioral performance across syllables suggest that the characteristics of sound stimuli influence the difficulty of the task and may affect the behavioral advantages of having richer, more robust spatial cues (Kidd et al., 2010). Concordantly, a previous study with complex tone stimuli has shown much larger differences in behavioral performance, up to 20% (Schröger, 1996), whereas studies presenting speech stimuli in a multi-talker environment yielded no behavioral advantage of having combined cues compared to impoverished cues (Glyde et al., 2013). These behavioral discrepancies, in combination with our neural findings, indicate that behavioral performance alone is not a sensitive metric for determining whether cortical networks controlling spatial selective attention are fully engaged.

Non-individualized or generic HRTFs such as from another listener or a mannikin have also been used widely for sound spatialization in auditory neuroscience studies (e.g., Choi et al., 2013; Klatt et al., 2018; Warren and Griffiths, 2003). Early psychoacoustic investigations (Middlebrooks, 1999; Wenzel et al., 1993) as well as a more recent EEG study (Wisniewski et al., 2016) demonstrated large inter-individual differences in the deteriorating effect of using generic HRTFs on localization abilities, mainly along the up-down and front-back dimensions. Although these ad-hoc degradations are predictable based on spectral comparisons with the listener-specific HRTFs (Baumgartner et al., 2016, 2014), it is poorly understood why some listeners adapt much faster than others to generic HRTFs, also without providing explicit feedback (e.g., Stitt et al., 2019). Because our study was not targeted to investigate such inter-individual differences, we aimed to reduce inter-subject variability by individualized HRTFs and did not include a spatialization condition using generic HRTFs. If individual HRTF measurements are not feasible it is advisable to individually select HRTFs from a database (e.g., Stitt et al., 2019; Warren and Griffiths, 2003).

## Conclusions

Our results indicate that although impoverished spatial cues can support spatial segregation of speech in a multi-talker environment, they do not fully engage the brain networks controlling spatial attention and lead to weak attentional control. Previous auditory studies have provided evidence that impoverished spatial cues do not evoke the same neural processing mechanisms as natural cue combinations during localization tasks with single sounds (Callan et al., 2013; Getzmann and Lewald, 2010; Palomäki et al., 2005). The current study extends these findings, demonstrating that the efficacy of higher-level cognitive processing, such as deployment of auditory selective attention, also depends on the naturalness of spatial cues. Poor attentional control was reflected in limited modulation of neural biomarkers of attentional processes. These findings suggest that the many past auditory attention studies using impoverished spatial cues may have underestimated the robust changes in cortical activity associated with deployment of spatial auditory attention in natural settings. Although impoverished auditory spatial cues can allow listeners to deploy spatial attention effectively enough to perform well in simple acoustic scenes, noisy, complex listening environments like those encountered in everyday environments pose greater challenges to attentional processing. In natural settings, spatial attention may fail unless attentional control networks are fully engaged. Thus, these results demonstrate the importance of preserving rich, natural spatial cues in hearing aids and other assistive listening devices.

## Acknowledgements

We thank Ashvini Meltoke for collecting part of the data, Gerald Kidd Jr. for providing the facilities for individualized HRTF measurements, and Virginia Best for fruitful discussions. This work was supported by the Austrian Science Fund (grant J3803-N30 to R.B.), European Commission (grant 691229 to R.B.), and NIDCD (grant DC013825 to B.S.-C.).

## References

Ahveninen, J., Hämäläinen, M., Jääskeläinen, I.P., Ahlfors, S.P., Huang, S., Lin, F.-H., Raij, T., Sams, M., Vasios, C.E., Belliveau, J.W., 2011. Attention-driven auditory cortex short-term plasticity helps segregate relevant sounds from noise. Proceedings of the National Academy of Sciences 108, 4182–4187. https://doi.org/10.1073/pnas.1016134108

Alho, K., Medvedev, S.V., Pakhomov, S.V., Roudas, M.S., Tervaniemi, M., Reinikainen, K., Zeffiro, T., Näätänen, R., 1999a. Selective tuning of the left and right auditory cortices during spatially directed attention. Cognitive Brain Research 7, 335–341. https://doi.org/10.1016/S0926-6410(98)00036-6

Banerjee, S., Snyder, A.C., Molholm, S., Foxe, J.J., 2011. Oscillatory Alpha-Band Mechanisms and the Deployment of Spatial Attention to Anticipated Auditory and Visual Target Locations: Supramodal or Sensory-Specific Control Mechanisms? Journal of Neuroscience 31, 9923–9932. https://doi.org/10.1523/jneurosci.4660-10.2011

Baumgartner, R., Majdak, P., Laback, B., 2016. Modeling the Effects of Sensorineural Hearing Loss on Sound Localization in the Median Plane. Trends in Hearing 20, 1–11. https://doi.org/10.1177/2331216516662003

Baumgartner, R., Majdak, P., Laback, B., 2014. Modeling sound-source localization in sagittal planes for human listeners. The Journal of the Acoustical Society of America 136, 791–802. https://doi.org/10.1121/1.4887447

Baumgartner, R., Reed, D.K., Tóth, B., Best, V., Majdak, P., Colburn, H.S., Shinn-Cunningham, B., 2017. Asymmetries in behavioral and neural responses to spectral cues demonstrate the generality of auditory looming bias. Proceedings of the National Academy of Sciences 114, 9743–9748. https://doi.org/10.1073/pnas.1703247114

Blankertz, B., Lemm, S., Treder, M., Haufe, S., Müller, K.-R., 2011. Single-trial analysis and classification of ERP components — A tutorial. NeuroImage 56, 814–825. https://doi.org/10.1016/J.NEUROIMAGE.2010.06.048

Blauert, J., 1997b. Spatial hearing. The Psychophysics of Human Sound Localization, 2nd edition. ed. MIT-Press, Cambridge, MA.

Boersma, P., 2001. Praat, a system for doing phonetics by computer. Glot International 5, 341–345.

Brainard, D.H., 1997. The psychophysics toolbox. Spatial vision 10, 433–436.

Callan, A., Callan, D.E., Ando, H., 2013. Neural correlates of sound externalization. NeuroImage 66, 22–27. https://doi.org/10.1016/j.neuroimage.2012.10.057

Chait, M., de Cheveigné, A., Poeppel, D., Simon, J.Z., 2010. Neural dynamics of attending and ignoring in human auditory cortex. Neuropsychologia 48, 3262–3271. https://doi.org/10.1016/J.NEUROPSYCHOLOGIA.2010.07.007

Cherry, E.C., 1953. Some Experiments on the Recognition of Speech, with One and with Two Ears. The Journal of the Acoustical Society of America 25, 975–979. https://doi.org/10.1121/1.1907229

Choi, I., Rajaram, S., Varghese, L.A., Shinn-Cunningham, B.G., 2013. Quantifying attentional modulation of auditory-evoked cortical responses from single-trial electroencephalography. Frontiers in Human Neuroscience 7, 115. https://doi.org/10.3389/fnhum.2013.00115

Couperus, J.W., Mangun, G.R., 2010. Signal enhancement and suppression during visual–spatial selective attention. Brain Research 1359, 155–177. https://doi.org/10.1016/j.brainres.2010.08.076

Cubick, J., Buchholz, J.M., Best, V., Lavandier, M., Dau, T., 2018. Listening through hearing aids affects spatial perception and speech intelligibility in normal-hearing listeners. The Journal of the Acoustical Society of America 144, 2896–2905. https://doi.org/10.1121/1.5078582

Culling, J.F., Hawley, M.L., Litovsky, R.Y., 2004. The role of head-induced interaural time and level differences in the speech reception threshold for multiple interfering sound sources. The Journal of the Acoustical Society of America 116, 1057–1065. https://doi.org/10.1121/1.1772396

Cusack, R., Carlyon, R.P., Robertson, I.H., 2001. Auditory Midline and Spatial Discrimination in Patients with Unilateral Neglect. Cortex 37, 706–709. https://doi.org/10.1016/S0010-9452(08)70620-8

Dahmen, J.C., Keating, P., Nodal, F.R., Schulz, A.L., King, A.J., 2010. Adaptation to Stimulus Statistics in the Perception and Neural Representation of Auditory Space. Neuron 66, 937–948. https://doi.org/10.1016/J.NEURON.2010.05.018

Dai, L., Best, V., Shinn-Cunningham, B.G., 2018. Sensorineural hearing loss degrades behavioral and physiological measures of human spatial selective auditory attention. Proceedings of the National Academy of Sciences 201721226. https://doi.org/10.1073/pnas.1721226115

Das, N., Biesmans, W., Bertrand, A., Francart, T., 2016. The effect of head-related filtering and ear-specific decoding bias on auditory attention detection. Journal of Neural Engineering 13, 056014. https://doi.org/10.1088/1741-2560/13/5/056014

Delorme, A., Makeig, S., 2004. EEGLAB: an open source toolbox for analysis of single-trial EEG dynamics including independent component analysis. Journal of Neuroscience Methods 134, 9–21. https://doi.org/10.1016/j.jneumeth.2003.10.009

Delorme, A., Sejnowski, T., Makeig, S., 2007. Enhanced detection of artifacts in EEG data using higher-order statistics and independent component analysis. NeuroImage 34, 1443–1449. https://doi.org/10.1016/j.neuroimage.2006.11.004

Deng, Y., Choi, I., Shinn-Cunningham, B., Baumgartner, R., 2019. Impoverished auditory cues limit engagement of brain networks controlling spatial selective attention. https://doi.org/10.5281/zenodo.3357164

Ellinger, R.L., Jakien, K.M., Gallun, F.J., 2017. The role of interaural differences on speech intelligibility in complex multi-talker environmentsa). The Journal of the Acoustical Society of America 141, EL170–EL176. https://doi.org/10.1121/1.4976113

Foxe, J.J., Snyder, A.C., 2011. The Role of Alpha-Band Brain Oscillations as a Sensory Suppression Mechanism during Selective Attention. Frontiers in Psychology 2, 154. https://doi.org/10.3389/fpsyg.2011.00154

Freunberger, R., Höller, Y., Griesmayr, B., Gruber, W., Sauseng, P., Klimesch, W., 2008. Functional similarities between the P1 component and alpha oscillations. European Journal of Neuroscience 27, 2330–2340. https://doi.org/10.1111/j.1460-9568.2008.06190.x

Fukuda, K., Vogel, E.K., 2009. Human Variation in Overriding Attentional Capture. Journal of Neuroscience 29, 8726–8733. https://doi.org/10.1523/jneurosci.2145-09.2009

Getzmann, S., Lewald, J., 2010. Effects of natural versus artificial spatial cues on electrophysiological correlates of auditory motion. Hearing Research 259, 44–54. https://doi.org/10.1016/j.heares.2009.09.021

Giuliano, R.J., Karns, C.M., Neville, H.J., Hillyard, S.A., 2014. Early Auditory Evoked Potential Is Modulated by Selective Attention and Related to Individual Differences in Visual Working Memory Capacity. Journal of Cognitive Neuroscience. https://doi.org/10.1162/jocn_a_00684

Glyde, H., Buchholz, J.M., Dillon, H., Cameron, S., Hickson, L., 2013. The importance of interaural time differences and level differences in spatial release from masking. The Journal of the Acoustical Society of America 134, EL147–EL152. https://doi.org/10.1121/1.4812441

Haegens, S., Handel, B.F., Jensen, O., 2011. Top-Down Controlled Alpha Band Activity in Somatosensory Areas Determines Behavioral Performance in a Discrimination Task. Journal of Neuroscience 31, 5197–5204. https://doi.org/10.1523/jneurosci.5199-10.2011

Hartmann, W.M., Wittenberg, A., 1996. On the externalization of sound images. J Acoust Soc Am 99, 3678–88.

Herrmann, C.S., Knight, R.T., 2001. Mechanisms of human attention: event-related potentials and oscillations. Neuroscience & Biobehavioral Reviews 25, 465–476. https://doi.org/10.1016/S0149-7634(01)00027-6

Higgins, N.C., McLaughlin, S.A., Rinne, T., Stecker, G.C., 2017. Evidence for cue-independent spatial representation in the human auditory cortex during active listening. Proceedings of the National Academy of Sciences 114, E7602--E7611. https://doi.org/10.1073/pnas.1707522114

Hillyard, S.A., Anllo-Vento, L., 1998. Event-related brain potentials in the study of visual selective attention. Proceedings of the National Academy of Sciences 95, 781–787. https://doi.org/10.1073/pnas.95.3.781

Hillyard, S.A., Vogel, E.K., Luck, S.J., 1998. Sensory gain control (amplification) as a mechanism of selective attention: electrophysiological and neuroimaging evidence. Philosophical transactions of the Royal Society of London. Series B, Biological sciences 353, 1257–70. https://doi.org/10.1098/rstb.1998.0281

Hopfinger, J., Luck, S., Hillyard, S., 2004. Selective attention: Electrophysiological and neuromagnetic studies. The cognitive neurosciences III 561–574.

Itoh, K., Yumoto, M., Uno, A., Kurauchi, T., Kaga, K., 2000. Temporal stream of cortical representation for auditory spatial localization in human hemispheres. Neuroscience Letters 292, 215–219. https://doi.org/10.1016/S0304-3940(00)01465-8

Jensen, O., Mazaheri, A., 2010. Shaping Functional Architecture by Oscillatory Alpha Activity: Gating by Inhibition. Frontiers in Human Neuroscience 4, 186. https://doi.org/10.3389/fnhum.2010.00186

Johnson, B.W., Hautus, M.J., 2010. Processing of binaural spatial information in human auditory cortex: {Neuromagnetic} responses to interaural timing and level differences. Neuropsychologia 48, 2610–2619. https://doi.org/10.1016/J.NEUROPSYCHOLOGIA.2010.05.008

Kelly, S.P., 2006. Increases in Alpha Oscillatory Power Reflect an Active Retinotopic Mechanism for Distracter Suppression During Sustained Visuospatial Attention. Journal of Neurophysiology 95, 3844–3851. https://doi.org/10.1152/jn.01234.2005

Kidd, G., Mason, C.R., Best, V., Marrone, N., Marrone, N., 2010. Stimulus factors influencing spatial release from speech-on-speech masking. The Journal of the Acoustical Society of America 128, 1965–78. https://doi.org/10.1121/1.3478781

Klatt, L.-I., Getzmann, S., Wascher, E., Schneider, D., 2018. The contribution of selective spatial attention to sound detection and sound localization: Evidence from event-related potentials and lateralized alpha oscillations. Biological Psychology 138, 133–145. https://doi.org/10.1016/j.biopsycho.2018.08.019

Klimesch, W., 2012. Alpha-band oscillations, attention, and controlled access to stored information. Trends in Cognitive Sciences 16, 606–617. https://doi.org/10.1016/J.TICS.2012.10.007

Klimesch, W., 2011. Evoked alpha and early access to the knowledge system: The P1 inhibition timing hypothesis. Brain Research. https://doi.org/10.1016/j.brainres.2011.06.003

Klimesch, W., Sauseng, P., Hanslmayr, S., 2007. EEG alpha oscillations: The inhibition-timing hypothesis. Brain Research Reviews 53, 63–88. https://doi.org/10.1016/j.brainresrev.2006.06.003

Kong, L., Michalka, S.W., Rosen, M.L., Sheremata, S.L., Swisher, J.D., Shinn-Cunningham, B.G., Somers, D.C., 2014. Auditory Spatial Attention Representations in the Human Cerebral Cortex. Cerebral Cortex 24, 773–784. https://doi.org/10.1093/cercor/bhs359

Lim, S.-J., Wöstmann, M., Obleser, J., 2015. Selective Attention to Auditory Memory Neurally Enhances Perceptual Precision. Journal of Neuroscience 35, 16094–16104. https://doi.org/10.1523/JNEUROSCI.2674-15.2015

Loiselle, L.H., Dorman, M.F., Yost, W.A., Cook, S.J., Gifford, R.H., 2016. Using ILD or ITD Cues for Sound Source Localization and Speech Understanding in a Complex Listening Environment by Listeners With Bilateral and With Hearing-Preservation Cochlear Implants. Journal of speech, language, and hearing research: JSLHR 59, 810–8. https://doi.org/10.1044/2015_JSLHR-H-14-0355

Luck, S.J., 1995. Multiple mechanisms of visual-spatial attention: recent evidence from human electrophysiology. Behavioural Brain Research 71, 113–123. https://doi.org/10.1016/0166-4328(95)00041-0

Majdak, P., Baumgartner, R., Jenny, C., 2019. Formation of three-dimensional auditory space. 1901.03990 [q-bio].

Marzecová, A., Schettino, A., Widmann, A., SanMiguel, I., Kotz, S.A., Schröger, E., 2018. Attentional gain is modulated by probabilistic feature expectations in a spatial cueing task: ERP evidence. Scientific Reports 8. https://doi.org/10.1038/s41598-017-18347-1

Mesgarani, N., Chang, E.F., 2012. Selective cortical representation of attended speaker in multi-talker speech perception. Nature 485, 233–236. https://doi.org/10.1038/nature11020

Middlebrooks, J.C., 2015. Sound localization. Handbook of Clinical Neurology, The Human Auditory System 129, 99–116. https://doi.org/10.1016/B978-0-444-62630-1.00006-8

Middlebrooks, J.C., 1999. Individual differences in external-ear transfer functions reduced by scaling in frequency. J. Acoust. Soc. Am. 106 (3), 1480–1492.

Murray, M.M., Brunet, D., Michel, C.M., 2008. Topographic ERP Analyses: A Step-by-Step Tutorial Review. Brain Topography 20, 249–264. https://doi.org/10.1007/s10548-008-0054-5

Palomäki, K.J., Tiitinen, H., Mäkinen, V., May, P.J.C., Alku, P., 2005. Spatial processing in human auditory cortex: The effects of 3D, ITD, and ILD stimulation techniques. Cognitive Brain Research 24, 364–379. https://doi.org/10.1016/j.cogbrainres.2005.02.013

Posner, M.I., Snyder, C.R., Davidson, B.J., 1980. Attention and the detection of signals. Journal of Experimental Psychology: General 109, 160–174. https://doi.org/10.1037/0096-3445.109.2.160

Rihs, T.A., Michel, C.M., Thut, G., 2007. Mechanisms of selective inhibition in visual spatial attention are indexed by ?-band EEG synchronization. European Journal of Neuroscience 25, 603–610. https://doi.org/10.1111/j.1460-9568.2007.05278.x

Rimmele, J.M., Zion Golumbic, E., Schröger, E., Poeppel, D., 2015. The effects of selective attention and speech acoustics on neural speech-tracking in a multi-talker scene. Cortex 68, 144–154. https://doi.org/10.1016/J.CORTEX.2014.12.014

Röder, B., Teder-Sälejärvi, W., Sterr, A., Rösler, F., Hillyard, S.A., Neville, H.J., 1999. Improved auditory spatial tuning in blind humans. Nature 400, 162–166. https://doi.org/10.1038/22106

Sach, A.J., Hill, N.I., Bailey, P.J., 2000. Auditory spatial attention using interaural time differences. Journal of Experimental Psychology: Human Perception and Performance 26, 717–729. https://doi.org/10.1037/0096-1523.26.2.717

Salminen, N.H., Takanen, M., Santala, O., Lamminsalo, J., Altoè, A., Pulkki, V., 2015. Integrated processing of spatial cues in human auditory cortex. Hearing Research 327, 143–152. https://doi.org/10.1016/J.HEARES.2015.06.006

Samaha, J., Bauer, P., Cimaroli, S., Postle, B.R., 2015. Top-down control of the phase of alpha-band oscillations as a mechanism for temporal prediction. Proceedings of the National Academy of Sciences of the United States of America 112, 8439–8444. https://doi.org/10.1073/pnas.1503686112

Samaha, J., Sprague, T.C., Postle, B.R., 2016. Decoding and Reconstructing the Focus of Spatial Attention from the Topography of Alpha-band Oscillations. Journal of Cognitive Neuroscience 28, 1090–1097. https://doi.org/10.1162/jocn_a_00955

Sauseng, P., Klimesch, W., Doppelmayr, M., Pecherstorfer, T., Freunberger, R., Hanslmayr, S., 2005. EEG alpha synchronization and functional coupling during top-down processing in a working memory task. Human Brain Mapping 26, 148–155. https://doi.org/10.1002/hbm.20150

Schreuder, M., Rost, T., Tangermann, M., 2011. Listen, You are Writing! Speeding up Online Spelling with a Dynamic Auditory BCI. Frontiers in Neuroscience 5, 112. https://doi.org/10.3389/fnins.2011.00112

Schröger, E., 1996. Interaural time and level differences: {Integrated} or separated processing? Hearing Research 96, 191–198. https://doi.org/10.1016/0378-5955(96)00066-4

Shinn-Cunningham, B.G., Kopco, N., Martin, T.J., 2005. Localizing nearby sound sources in a classroom: Binaural room impulse responses. The Journal of the Acoustical Society of America 117, 3100–3115. https://doi.org/10.1121/1.1872572

Siegel, M., Donner, T.H., Engel, A.K., 2012. Spectral fingerprints of large-scale neuronal interactions. Nature Reviews Neuroscience 13, 121–134. https://doi.org/10.1038/nrn3137

Skrandies, W., 1990. Global field power and topographic similarity. Brain Topography 3, 137–141.

Snyder, J.S., Large, E.W., 2005. Gamma-band activity reflects the metric structure of rhythmic tone sequences. Cognitive Brain Research 24, 117–126. https://doi.org/10.1016/j.cogbrainres.2004.12.014

Stevens, C., Fanning, J., Coch, D., Sanders, L., Neville, H., 2008. Neural mechanisms of selective auditory attention are enhanced by computerized training: Electrophysiological evidence from language-impaired and typically developing children. Brain Research 1205, 55–69. https://doi.org/10.1016/J.BRAINRES.2007.10.108

Stitt, P., Picinali, L., Katz, B.F.G., 2019. Auditory Accommodation to Poorly Matched Non-Individual Spectral Localization Cues Through Active Learning. Scientific Reports 9. https://doi.org/10.1038/s41598-018-37873-0

Tardif, E., Murray, M.M., Meylan, R., Spierer, L., Clarke, S., 2006. The spatio-temporal brain dynamics of processing and integrating sound localization cues in humans. Brain Research 1092, 161–176. https://doi.org/10.1016/J.BRAINRES.2006.03.095

Ungan, P., Yagcioglu, S., Goksoy, C., 2001. Differences between the N1 waves of the responses to interaural time and intensity disparities: scalp topography and dipole sources. Clinical Neurophysiology 112, 485–498. https://doi.org/10.1016/S1388-2457(00)00550-2

Warren, J.D., Griffiths, T.D., 2003. Distinct Mechanisms for Processing Spatial Sequences and Pitch Sequences in the Human Auditory Brain. J. Neurosci. 23, 5799–5804. https://doi.org/10.1523/JNEUROSCI.23-13-05799.2003

Wenzel, E.M., Arruda, M., Kistler, D.J., Wightman, F.L., 1993. Localization using nonindividualized head-related transfer functions. The Journal of the Acoustical Society of America 94, 111–123. https://doi.org/10.1121/1.407089

Wisniewski, M.G., Romigh, G.D., Kenzig, S.M., Iyer, N., Thompson, E.R., Simpson, B.D., Rothwell, C.D., 2016. Enhanced auditory spatial performance using individualized head-related transfer functions: An event-related potential study. The Journal of the Acoustical Society of America 139, 1993–1994. https://doi.org/10.1121/1.4949839

Worden, M.S., Foxe, J.J., Wang, N., Simpson, G. V., 2018. Anticipatory Biasing of Visuospatial Attention Indexed by Retinotopically Specific α-Bank Electroencephalography Increases over Occipital Cortex. The Journal of Neuroscience 20, RC63–RC63. https://doi.org/10.1523/jneurosci.20-06-j0002.2000

Wöstmann, M., Herrmann, B., Maess, B., Obleser, J., 2016. Spatiotemporal dynamics of auditory attention synchronize with speech. Proceedings of the National Academy of Sciences 113, 3873–3878. https://doi.org/10.1073/pnas.1523357113

Wyart, V., Dehaene, S., Tallon-Baudry, C., 2012. Early dissociation between neural signatures of endogenous spatial attention and perceptual awareness during visual masking. Frontiers in Human Neuroscience 6, 16. https://doi.org/10.3389/fnhum.2012.00016

